# Intra-tumor heterogeneity defines treatment-resistant HER2+ breast tumors

**DOI:** 10.1101/297549

**Authors:** Inga H. Rye, Anne Trinh, Anna Sætersdal, Daniel Nebdal, Ole Christian Lingjærde, Vanessa Almendro, Kornelia Polyak, Anne-Lise Børresen-Dale, Åslaug Helland, Florian Markowetz, Hege G. Russnes

**Author notes:** Equal contribution (Inga H. Rye, Anne Trinh). Department of Medical Oncology, Dana-Farber Cancer Institute, Boston (Anne Trinh). Vertex pharmaceuticals, Boston (Vanessa Almendro). Shared last author (Florian Markowetz, Hege G. Russnes). Corresponding author: Hege G. Russnes, Department of Cancer Genetics, Institute for Cancer Research, Oslo University Hospital Radiumhospitalet, 0424 Oslo, Norway.

## Abstract

Targeted therapy for patients with HER2 positive (HER2+) breast cancer has improved the overall survival, but many patients still suffer relapse and death of the disease. Intra-tumor heterogeneity of both estrogen receptor (ER) and HER2 expression has been proposed to play a key role in treatment failure, but little work has been done to comprehensively study this heterogeneity at the single-cell level.

In this study, we explored the clinical impact of intra-tumor heterogeneity of ER protein expression, HER2 protein expression, and *HER2* gene copy number alterations. Using combined immunofluorescence and *in situ* hybridization on tissue sections followed by a validated computational approach, we analyzed more than 13,000 single tumor cells across 37 HER2+ breast tumors. The samples were taken both before and after neoadjuvant chemotherapy plus HER2-targeted treatment, enabling us to study tumor evolution as well.

We found that intra-tumor heterogeneity for *HER2* copy number varied substantially between patient samples. Highly heterogeneous tumors were associated with significantly shorter disease-free survival and fewer long-term survivors. Patients for which *HER2* characteristics did not change during treatment had a significantly worse outcome.

This work shows the impact of intra-tumor heterogeneity in molecular diagnostics for treatment selection in HER2+ breast cancer patients and the power of computational scoring methods to evaluate *in situ* molecular markers in tissue biopsies.

## 1 Introduction

Breast cancer is divided into several distinct subtypes and the expression level of estrogen receptor (ER), progesterone receptor (PgR) and human epidermal growth factor receptor 2 (HER2) are fundamental for treatment decision and prognosis of the disease. The HER2 positive (HER2+) tumors account for 15-20% of all breast cancers and are characterized by either over-expression of HER2 protein and/or increased copy number of the *HER2* gene. With the introduction of HER2-targeted therapy, such as trastuzumab and lapatinib, the overall survival for both early and late stage disease has increased (Baselga et al. 2012; Cortazar et al. 2014; Gianni et al. 2010; Guarneri and Conte 2004; Viani et al. 2007).

Breast cancer was one of the first solid cancer types where comprehensive molecular profiling revealed robust molecular subtypes (Curtis et al. 2012; Perou et al. 2000), and HER2+ tumors are found within several subtypes. By PAM50 classification, HER2+ tumors are mainly found in the HER2-enriched but also in the luminal B and luminal A subtypes (Parker et al. 2009). Similarly, in the 10 integrated cluster (IntClust) subtypes, the HER2+ tumors dominate group 5 but are also found within other subtypes (Curtis et al. 2012). The notion that HER2+ tumors do not represent a separate subtype but a wider biological spectrum was strengthened by a recent study identifying four different subtypes of HER2+ breast carcinomas based on gene expression signatures (Ferrari et al. 2016).

Pathologists have noticed the presence of cell-to-cell variation in HER2+ tumors since the introduction of biomarkers into diagnostic routine. In early stage HER2+ breast cancer, neither the average level of HER2 protein expression nor the average level of *HER2* gene amplification across a tumor seem to have an impact on therapy response (Wolff et al. 2013; Zabaglo et al. 2013). However, as reflected by the comprehensive College of American Pathologists (CAP) guidelines, some HER2+ tumors display intra-tumor variation in *HER2* copy number (*HER2* CN) levels. The ASCO/CAP guidelines from 2013 state that breast cancers with aggregations of *HER2* amplified cells (with *HER2*/CEP17 ratio >2.0 or more than 6 *HER2* copies per cell) in more than 10% of the tumor must be quantified and reported separately (Wolff et al. 2013). The clinical challenge of such a definition has been addressed for HER2 equivocal cases (Bartlett et al. 2011; Lewis et al. 2005), but the clinical impact of intra-tumor heterogeneity within non-equivocal HER2+ tumors are less studied (Arena et al. 2013; Gulbahce et al. 2016). The regional variation of *HER2* gene amplification has been studied to some extent (Lee et al. 2014; Seol et al. 2012) and heterogeneity of *HER2* CN even in tumors classified as non-amplified was recently described (Buckley et al. 2016), but there are very few studies addressing this at the single cell level estimating multiple biomarkers from a high number of cells.

To investigate and quantify the heterogeneity of HER2+ carcinomas by using single cell investigation, we performed detailed *in situ* analyses on samples from a Norwegian observational study (RA-HER2), comprised of 37 HER2+ patients treated in a neoadjuvant setting with trastuzumab and chemotherapy where both response data as well as clinical follow up were available. For objective assessment of the molecular *in situ* markers we used GoIFISH, a software for image analysis developed to objectively score both immunofluorescence and FISH signals from numerous individual tumors cells (Trinh et al. 2014). With this quantitative approach we examined 103 images and more than 13,000 cells showing the clinical impact of different types of genomic and phenotypic intra-tumor heterogeneity in HER2+ breast cancer.

## 2 Material and methods

### 2.1 Patient samples

Breast cancer patients diagnosed with HER2+ tumors between 2004-2010 who qualified for neoadjuvant treatment according to the national guidelines were included in this prospective observational trial. Informed consent was obtained from all patients, and the study was approved by the Regional Ethical Committee (South-east of Norway, no. S-06495b). The clinical characteristics are shown in Supplemental Table 1. All 37 patients received combinatorial neoadjuvant treatment of 4 cycles of fluorouracil, epirubicin and cyclophosphamide (FEC) followed by 4 cycles of taxanes in combination with the HER2 targeted monoclonal antibody trastuzumab. The average neoadjuvant treatment period was 6 months (range 3-10 months). The Response Evaluation Criteria In Solid Tumors (RECIST) (Nishino et al. 2010) was used to score the effect of the neoadjuvant treatment, with pathological complete response (pCR) defined as no invasive tumor cells in primary tumor region or lymph nodes after neoadjuvant treatment. Non-pCR was defined as presence of residual invasive tumor cells in primary tumor region or lymph nodes (Supplemental Table 1). After neoadjuvant treatment, 12 patients had pathological complete response (pCR), and among the 25 patients with non-complete pathological response (non-pCR), a variation in tumor reduction from almost complete response to no reduction in tumor size was observed (Supplemental Table 1).

Formalin-fixated paraffin-embedded (FFPE) tumor tissue from the 37 patients was collected from several hospitals throughout Norway. FFPE core needle biopsies from the time of diagnosis and FFPE surgical biopsies after neoadjuvant treatment were available for analysis. In addition, FFPE tissue biopsies from later distant metastases were available for 3 patients.

### 2.2 IFISH analyses

The FISH probes for *HER2* were made from the BAC clones RP11-94L15 and RP11-909L6, and FISH probes for centromere 17 (cent17) were made from BAC clones RP11-170N19 and RP11-909L10. The BAC probes were isolated according to the instructions from the manufacturer and labeled with fluorescent UTPS by nick translation. Primary antibody recognizing estrogen receptor (clone 6G11) were detected with secondary antibody IgG conjugated Alexa fluor 594. The HER2 (CB11) primary antibody was detected with a secondary biotinylated antibody and visualized using streptavidin conjugated Alexa fluor 488 antibody in order to visualize the protein expression of ER and HER2. A detailed IFISH protocol including antibody and BAC catalogue numbers is described in the previous publication (Trinh et al. 2014). The tissue samples were mounted with DAPI counterstain and areas of interest were photographed with 25 z-stacks in a Zeiss Axiovision M1 microscope. The areas with a high number of tumor cells and with high quality of IFISH staining were selected for photography. The number of biopsies, areas and tumor cells analyzed per sample are listed in Supplemental Table 2.

### 2.3 Analysis by GoIFISH

We previously developed and validated the software GoIFISH (Trinh et al. 2014), an image analysis pipeline designed to objectively recognize cell types, score protein intensities in distinct cellular compartments (nucleus, cytoplasm, and membranes), count and measure FISH spots/areas and intensities, measure nuclear size and display topological distributions of the cells and the analyzed parameters. GoIFISH estimates are highly concordant with visual scoring at the single cell level, and optimal intensity thresholds of 300 and 50 following adjustment by background and perinuclear intensity were used to define HER2 positive and ER positive cells respectively from 12-bit images. (Trinh et al. 2014). ER+ patients were identified according to the national guidelines with a cut-off level at 1% positive cells (Helsedirektoratet 2014). The *HER2* copy number (*HER2* CN) level was assessed by measuring the total area of the HER2 probe signals within each nucleus. For cluster analyses to study phenotypic heterogeneity we assigned each cell within a tumor into one of four phenotypic groups (HER2+/ER+, HER2+/ER-, HER2-/ER+, HER2-/ER-) based on the defined thresholds. To address heterogeneity based on genomic changes we assigned each cell into one of three *HER2* CN categories: normal (*HER2*norm), gain (*HER2*gain) or amplified (*HER2*amp). *HER2*norm reflected cells with up to 3 spots (0-63 pixels), *HER2*gain: 3-6 spots (64-200 pixels) and *HER2*amp: >6 spots (>200 pixels). Additionally we considered the combined effect of both phenotype and genotype and classified each cell into one of twelve groups: HER2+/ER+ *HER2* amp, HER2+/ER+ *HER2* gain, HER2+/ER+ *HER2* norm, HER2+/ER-*HER2* amp, HER2+/ER-*HER2* gain, HER2+/ER-*HER2* norm, HER2-/ER+ *HER2* amp, HER2-/ER+ *HER2* gain, HER2-/ER+ *HER2* norm, HER2-/ER-*HER2* amp, HER2-/ER-*HER2* gain or HER2-/ER-*HER2* norm.

Five samples were excluded in comparisons between pre- and post-treatment samples: three due to low numbers of tumor cells present after neoadjuvant therapy, and two samples had insufficient IFISH staining due to technical problems (immunofluorescence and genomic (FISH) analyses were performed separately).

### 2.4 Spatial distribution of HER2 amplification within tumor nuclei

Three spatial patterns of *HER2* FISH signals within individual tumor cell nuclei were identified by visual inspection. Cells demonstrating a tight cluster of multiple signals were called “cluster”, cells with distinct and separate signals were called “scatter” and those with both patterns were annotated as “mix”. The *HER2* spatial distribution pattern was scored in 100 tumor cells from each biopsy (from both pre- and post-treatment samples) and in the three samples from metastases. These single cell scores were collapsed to the patient level by (i) computing the frequency of each pattern and (ii) using a 70% majority cut-off to describe a class for each patient. If a tumor did not show one particular dominant pattern, it was considered as “heterogeneous”. In the pre-treatment samples, 10 were dominated by “cluster” cells, 6 with “mix”, 8 with “scatter” and 13 samples were “heterogeneous” with regard to spatial patterns.

### 2.5 Statistical analyses

The Welch t-test was used to determine differences in intensity distributions, Fisher exact t-test was used to calculate the differences between groups of patients. Survival curves were constructed using the Kaplan Meier method, using both disease free survival (i.e. time to metastasis) and overall breast cancer specific survival as events. Differences in survival between groups of patients were studied by univariate cox regression analyses and expressed as hazards ratios with 95% confidence intervals using continuous variables (corrected for age, stage and grade). The Shannon Index (SI) was used as measure for heterogeneity of the defined phenotypic, genomic groups and combined phenotypic and genomic groups (Shannon 1948), and the mean Shannon Index for each cluster group was used to determine the differences in heterogeneity between clusters.

To measure the change in the clonal composition during neoadjuvant therapy, the Kullback-Leibler divergence index (K-L) (Kullback and Leibler 1951) was used to compare the cell type distributions before and after treatment. Briefly, this describes the divergence between two populations, such as the phenotypic composition of pre- and post-treatment samples:

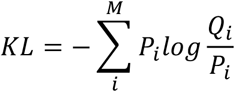

Where *P_i_* is the proportion of cells which belong to group in the pre-treatment group, and Q_i_ is the proportion of cells which belong to group i in the post-treatment samples. M indicates the number of discrete groups considered: four for phenotypic change, three for genomic changes and twelve for the combined change. A high index signifies different clonal compositions in the samples taken after treatment versus the samples taken before. The median of the Kullback-Leibler index was used to divide the samples into two equal sized groups, one group with samples with a high change of *HER2* CN fractions (K-L high) and one group with samples with low change in fractions (K-L low). All image analysis was performed in MATLAB (7.12.0(R2011a)), and subsequent statistical analyses were performed in R (R Core Team 2017).

## 3 Results

We analyzed more than 13 000 single tumor cells from biopsies taken before treatment (n=37), after treatment (n=22) and metastases (n=3) from 37 HER2+ positive breast cancer patients. Single-cell metrics for HER2 and ER expression, *HER2* copy number and CEP17 copy number were evaluated. This enabled us to evaluate the heterogeneity of the markers both across tumors but also within the individual tumors at different time points, as illustrated in Figure 1A-D. As an example, images of pre- and post-treatment biopsies from patient 7588 show the protein- and FISH staining of the tumor cells. The GoIFISH software was used to visualize the spatial distribution of cells with different phenotypic and/or genotypic features, as shown in Figure 1E-F where each cell is pseudo-colored with regard to HER2 and ER protein expression. Changes in cell populations during therapy are evident; prior to therapy the tumor had both HER2+/ER+ and HER2+/ER-negative cells, while in the post-treatment tumor a new dominant population of HER2-/ER+ cells emerged. The phenotypic change during therapy is further illustrated in Figure 1G, where each dot represents a tumor cell and the color illustrates the phenotype. Furthermore, there was a substantial reduction of cells with high *HER2* CN after treatment, reflected in Figure 1G by the size of each dot.

**Figure 1:**
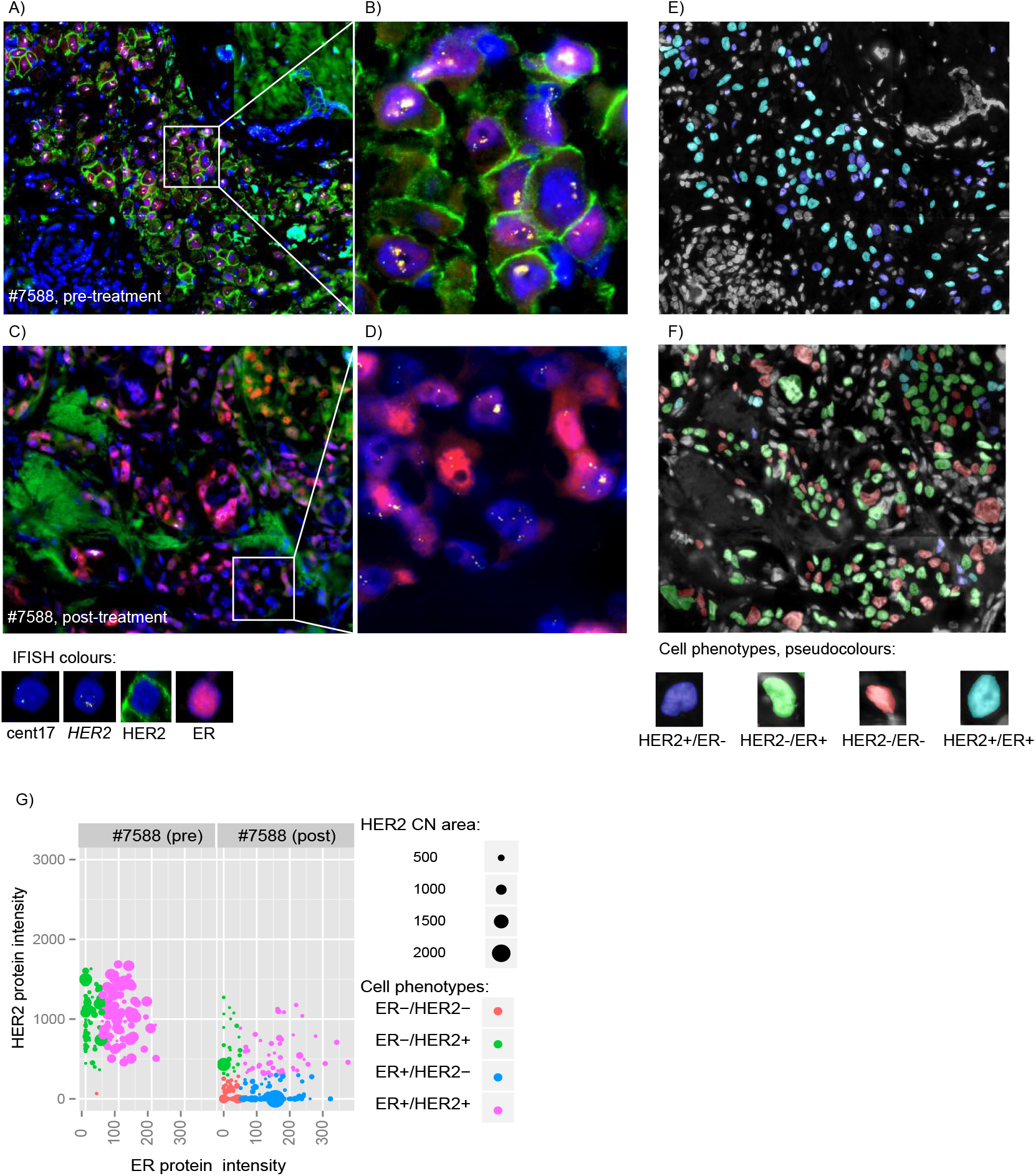
IFISH images reflecting intra tumor heterogeneity before and after treatment. Expression of ER and HER2 protein and copy number of *HER2* gene by IFISH (color code below images) for A) pre-treatment biopsy from patient #7588, B) magnified image of the outlined area, C) post-treatment biopsy of patient #7588 and D) magnified image of the outlined area. Pseudo-colored cell-phenotypes of E) pre-treatment biopsy (same area as in Figure 1A), F) post-treatment biopsy (same area as in Figure 1C). G) Tumor cell heterogeneity before and after treatment for patient #7588, the scatter plot shows the relationship between ER expression (y-axis) and HER2 expression (x-axis) for each of the individual cells. The color reflects the cell phenotype. The size of the dot reflects each cells *HER2* CN level, where a small dot equals fewer copies and a large dot more copies of the *HER2* gene.

### 3.1 Inter-tumor heterogeneity within HER2+ tumors

All images were subjected to the same analyses as for the case shown in Figure 1, and a substantial variation of marker distribution was seen across the cohort. This is visualized in the compilation of representative images from each of the 37 pre-treatment samples shown in Supplemental Figure 1. To get a first overview of the cohort, we estimated the mean values of the biomarkers (i.e. measurements from all tumor cells within a sample) and found patients with non-pathological complete response (non-pCR) to have a significant lower mean value of copy number of the *HER2* gene compared to patients with pathological complete response (pCR) (Supplemental Figure 2A, t-test: p = 0.02). No significant difference in mean HER2 and ER protein expression was found. By looking at the same biomarkers and stratifying the patients by disease progression we found a significant lower ER expression (p=0.02) and lower *HER2* CN/cent17 CN ratio (p=0.009) in samples from patients with later metastatic disease compared to those without metastasis (Figure 2A). Figure 2B illustrate the pre-treatment cell type composition in an ER negative tumor with highly amplified *HER2* CN from a patient which later had progressive disease. The cell composition in an ER positive tumor with gained *HER2* CN from a patient who has not had progressive disease is shown in Figure 2C.

**Figure 2:**
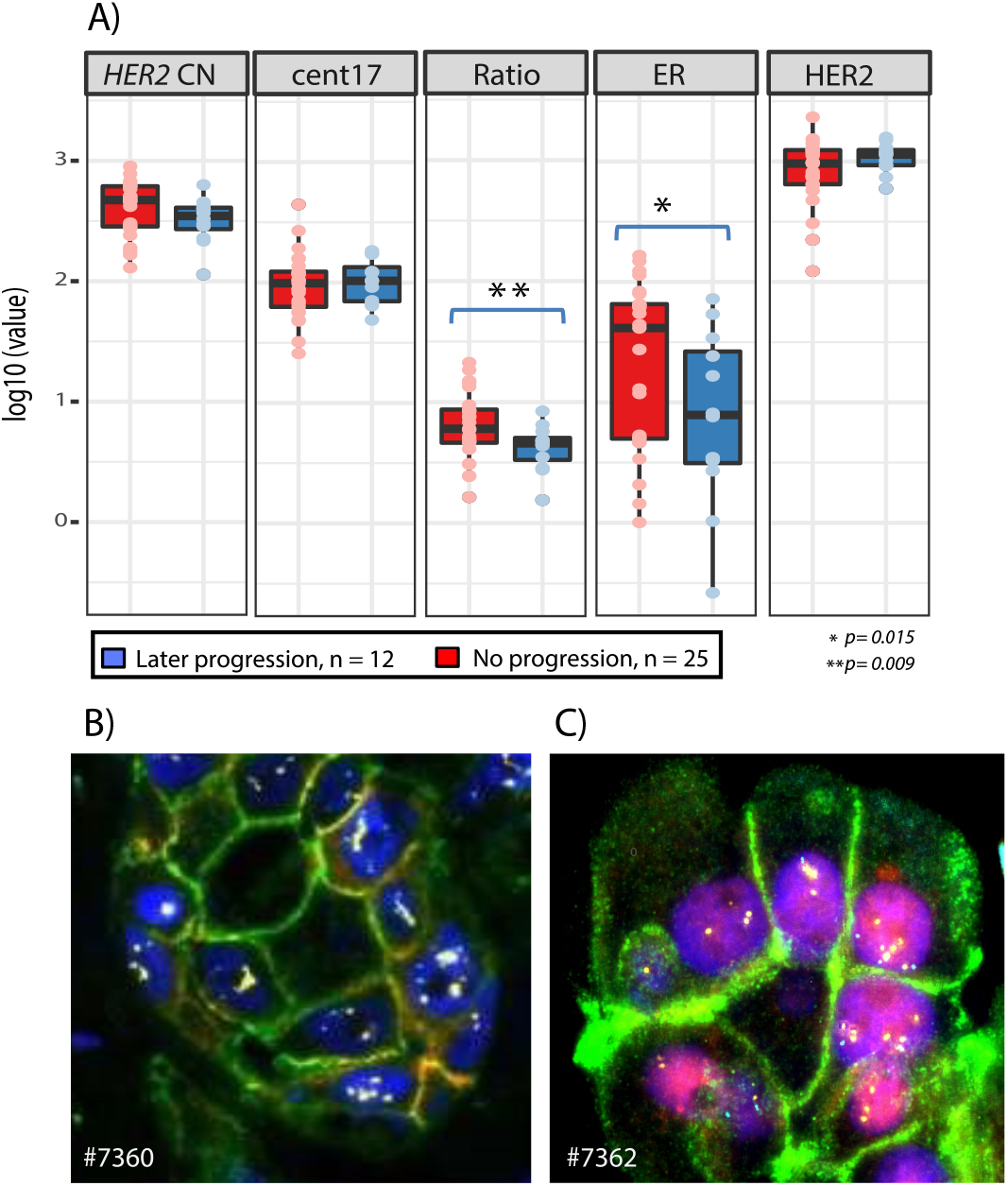
Biomarker status and later progression of disease. A) Comparison of GoIFISH measurements (*HER2* copy number (HER2 CN), cent17, ratio (HER2 CN/cent17), ER protein expression and HER2 protein expression) for all pre-treatment biopsies (n=37) stratified by relapse or not after neo-adjuvant treatment. B) IFISH image from a patient with later relapse of disease (#7360). The cells were ER-, HER2+ with amplification of *HER2* (same color scheme as in Figure 1 A-D). C) IFISH image from a patient without later relapse of the disease (#7362). The sample was ER+, HER2+ with gain of *HER2* copies.

Using 1% positive cells as a cut-off level from GoIFISH, we identified 28 patients with ER positive (ER+) tumors (76%) and nine patients with ER negative (ER-) tumors (24%). Complete response to neoadjuvant treatment was seen in 7/28 (28%) and 5/9 (55%) patients with ER+ and ER-tumors respectively. With regard to metastasis, 9/28 (32%) patients with ER+ and 3/9 (33%) patients with ER-tumors developed metastasis (Supplementary Table 1). Tumors were stratified into four groups based on the percentage of ER+ cells present: ER negative (<1%, n=9), low ER (1-10%, n=9), intermediate ER (10-50%, n=10) and high ER (>50%, n=9). Although not significant, a trend that patients with low or intermediate number of ER+ cells had less local response to treatment was observed, as well as a worse prognosis compared to those with either high ER or ER negative tumors (Supplementary Figure 2B).

We next sought to determine whether relationship between ER and HER2 protein expression and *HER2* copy number at a single-cell level could influence patient outcome. As illustrated by scatterplots in Supplemental Figure 3, a substantial variation was seen with regard to ER and HER2 protein expression both across tumors and within tumors. In addition, some tumors showed a linear relationship between *HER2* CN and HER2 protein level, but others did not (Supplemental Figure 4). In addition, we noticed that the relationship could change during therapy (Supplemental Figure 3 and 4).

To address the clinical implication of this protein variation, we assigned each cell to one of four categories; HER2+/ER+, HER2+/ER-, HER2-/ER+ or HER2-/ER- (see Methods section). By comparing the fractions of cells with different phenotypes, subsets of tumors with distinct types of phenotypic intra-tumor heterogeneity were identified. Hierarchical clustering of the fractions of each cell class revealed three separate groups of tumors. Group P1 contained tumors dominated by HER2+/ER+ cells while tumors in the cluster group P2 was dominated by HER2+/ER-cells (Figure 3A and Supplemental Table 3). IFISH images from two patients representing phenotypic cluster P1 and P2 are shown in Figure 3B. Patients in cluster group P2 had tumors with negative to intermediate ER expression and were associated with high histological grade (Supplemental Table 3). They also had a higher frequency of later metastasis, and the Kaplan Meier curves indicated a worse prognosis, although this was not significant (Figure 3C, Supplemental Figure 5A). Interestingly, P2 was the least heterogonous cluster with an Shannon index (SI) of 0.34, compared to P1 which had SI=0.66 (Supplemental Table 4). Cluster group P3 only contained three samples, all dominated by HER2 negative tumor cells. Two of these samples were scored 2+ by IHC (#7619 and #7441); the third sample (#7370) had one HER2 positive and one HER2 negative biopsy prior to therapy.

**Figure 3:**
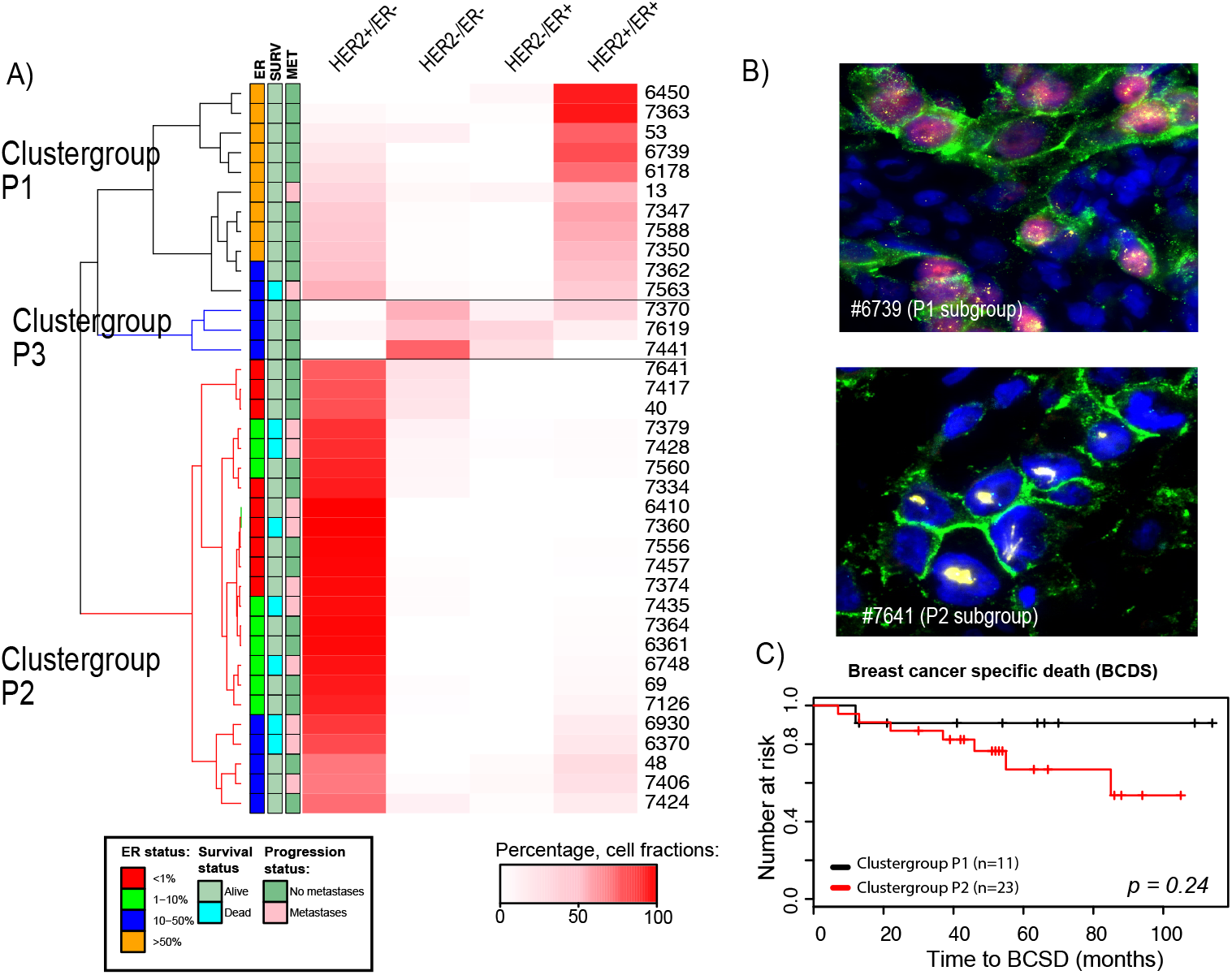
Identification of subsets of HER2+ breast cancer patients by phenotypic diversity. A) Unsupervised cluster analysis of the fractions of the phenotypic cell types HER2-/ER-, HER2+/ER-, HER2-/ER+ and HER2+/ER+ in the pre-treatment samples (n=37) where the percentage of each cell type (i.e. fraction) is indicated by the color intensity. Two large clusters and one small were identified, where clustergroup P1 (n=11) was dominated by HER2+/ER+ cells and clustergroup P2 was dominated by HER2+/ER-cells. The smallest clustergroup contained three patients whose tumors had mainly HER2-cells. The clinical information for each patient is illustrated by the boxes next to the dendrogram. B) IFISH image to the left is from pre-treatment biopsy from patient #6739 (in clustergroup P1) which was dominated by HER2+/ER+ tumor cells. The image to the right is from the pre-treatment sample from patient #7641 (clustergroup P2) dominated by HER2+/ER-tumor cells. C) Survival analyses; breast cancer specific death for the two groups (p=0.24). D) Survival analyses; breast cancer specific death between patients with different percentage of ER+ cells (p=0.14).

In contrast to cellular phenotypes, where subpopulations can be dynamic and cells might change expression levels rapidly in response to treatment, *HER2* copy number (CN) will reflect more persistent cellular subclones. We categorized each cell into one of three levels of *HER2* CN (*norm, gain* and *amp*), and determined the cellular composition of each tumor (see Methods section). We found some tumors to be dominated by cells with similar copy number level while other tumors had more heterogeneous cellular composition. Hierarchical clustering identified three groups of tumors with different levels of *HER2* genomic heterogeneity (Figure 4A, Supplemental Table 4) were identified. The most distinct difference between these three groups was the fraction of cells with *HER2* amplification. The smallest group of tumors (cluster group G1, n=6) had overall low level HER2 CN with few cells with *HER2*amp and the highest heterogeneity (SI=1.2). The second largest group (cluster group G2, n=13) had tumors mainly dominated by cells with *HER2*amp and had a low degree of heterogeneity (SI=0.6). This was in contrast to the third group (cluster group G3, n=16), which had a high fraction of *HER2*amp cells, but also fractions of *HER2*gain and *HER2*norm cells and overall a high degree of heterogeneity (SI=0.9). A representative image of cluster groups is shown in Figure 4B. Interestingly, the patients belonging to cluster G3 displaying high intra-tumor variation but with *HER2*amp dominating, were more likely to experience distant metastases (Supplemental Table 3) and had the highest risk of disease progression (HR: 14,9, p: 0.04, Figure 4C) but not a significant increased risk of death by breast cancer (Figure 4D). However, the groups were not distinguished by any other clinical parameter; we were in particular not able to find any significant correlation to treatment response measured by tumor reduction (Supplementary Table 3).

**Figure 4:**
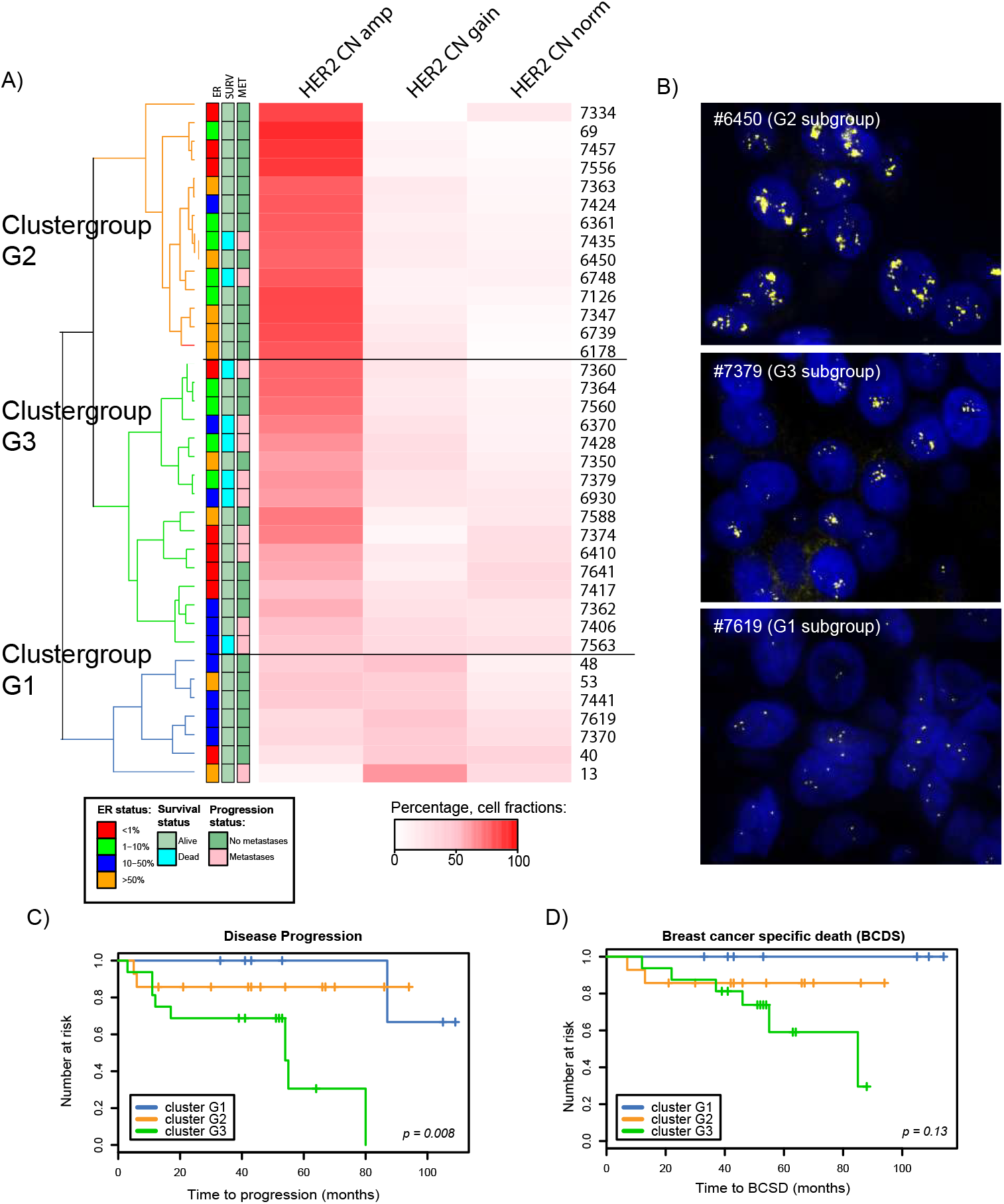
Identification of subsets of HER2+ breast cancer patients by HER2 copy number diversity. A) Unsupervised clustering based on the fractions of cells with different levels of *HER2* copy number (normal, gain or amplified). Three clusters (G1-G3) were identified. The clinical information for each patient is illustrated in the boxes next to the dendrogram. B) FISH (*HER2* CN) images from patient samples representing each of the three cluster groups (G1-G3). The top image is from cluster G2 (patient #6450) and shows a tumor dominated by *HER2* CN amp cell type, the second image is from cluster G3 (patient #7379) and shows a sample with an intermediate fraction of cells with *HER2* CN amp, the last image is from cluster G1 (#7619) and shows a sample with a high fraction of *HER2 CN* gain and a low fraction of *HER2* CN amp cell types. C) Survival analyses showed significant differences in risk for progression between the two groups (p=0.008) but not for breast cancer specific death D).

To investigate the impact of combined phenotypic and genomic heterogeneity, we next assigned each cell within a tumor to one of twelve combined phenotype-genomic (PG) groups (see Methods section). Three separate groups were identified (Supplementary Figure 5A), where cluster PG1 (n=9) was comprised of highly heterogeneous tumors containing both ER+ and ER-cells with varying *HER2* CN levels (amp, gain and norm) (SI=1.8). Cluster PG2 (n=8) consisted predominantly of tumors with ER+/HER2+ cells with *HER2*amp (SI=1.28). The largest group, cluster PG3 (n=20), was also dominated by cells with *HER2*amp with predominantly a ER-/HER2+ phenotype, but many tumors had cells with normal levels or gain of *HER2* CN (SI=0.99). Patients in PG2 had >50% ER+ cells and all had a high HER2 protein expression (3+) and none had later progression of the disease (Supplementary Table 3). Although not significant, a trend was observed where patients in the PG1 and PG3 groups had a higher risk for progressive disease and breast cancer related death than patients in group PG2 (Supplemental Figure 5B-C).

### 3.2 The HER2 spatial organization

During visual investigation of the images we noticed different spatial patterns of *HER2* amplifications within each nucleus. Some cells had a tight cluster of multiple signals, others had fewer signals scattered within the nucleus and some had a combination (Figure 5A, see Methods section for more details). We named the nuclear spatial patterns “cluster”, “scatter” and “mix”. As intra-tumor heterogeneity with regard to *HER2* CN levels seemed to have prognostic information, we wanted to address whether the observed differences in spatial organization of the HER2 gene was of clinical importance. As shown in the triangle plots in Figure 5B, we observed inter-tumor variation where some samples were dominated by one spatial type (samples in the corners of the triangle plot in Figure 5B) while other had a more heterogeneous distribution, illustrated by being plotted towards the centre of the triangle. A significant difference in the distribution of samples from patients with and without pathological complete response (pCR) was observed; samples from patients with pCR were most frequently of “cluster” or “mix” type while samples from patients with non-pCR were more heterogeneous and dominated the group characterized by the “scatter” type of distribution (Fisher’s exact test, p = 0.007, Supplemental Table 5A). We found an indication for patients with tumors dominated by “mixed” spatial type not to have disease progression, in contrast to patients with tumors dominated by “cluster” or with a combination of the three types (Figure 5C, Supplemental Figure 6A, Supplemental Table 5B). Interestingly, these spatial distributions were also associated with ER status: ER negative tumors were found to be frequently of “cluster” or “mix” spatial type (Supplemental Figure 6B, Supplemental Table 5C), and when stratifying the ER positive samples into negative, low (1-10%), intermediate (10-50%) and high ER (>50%), the intermediate ER+ tumors were predominantly of the “scatter” spatial type, while the ER negative and ER low tumors (p=0.007) were predominantly of the “cluster” spatial type. (Figure 5D, Supplemental Table 5D).

**Figure 5:**
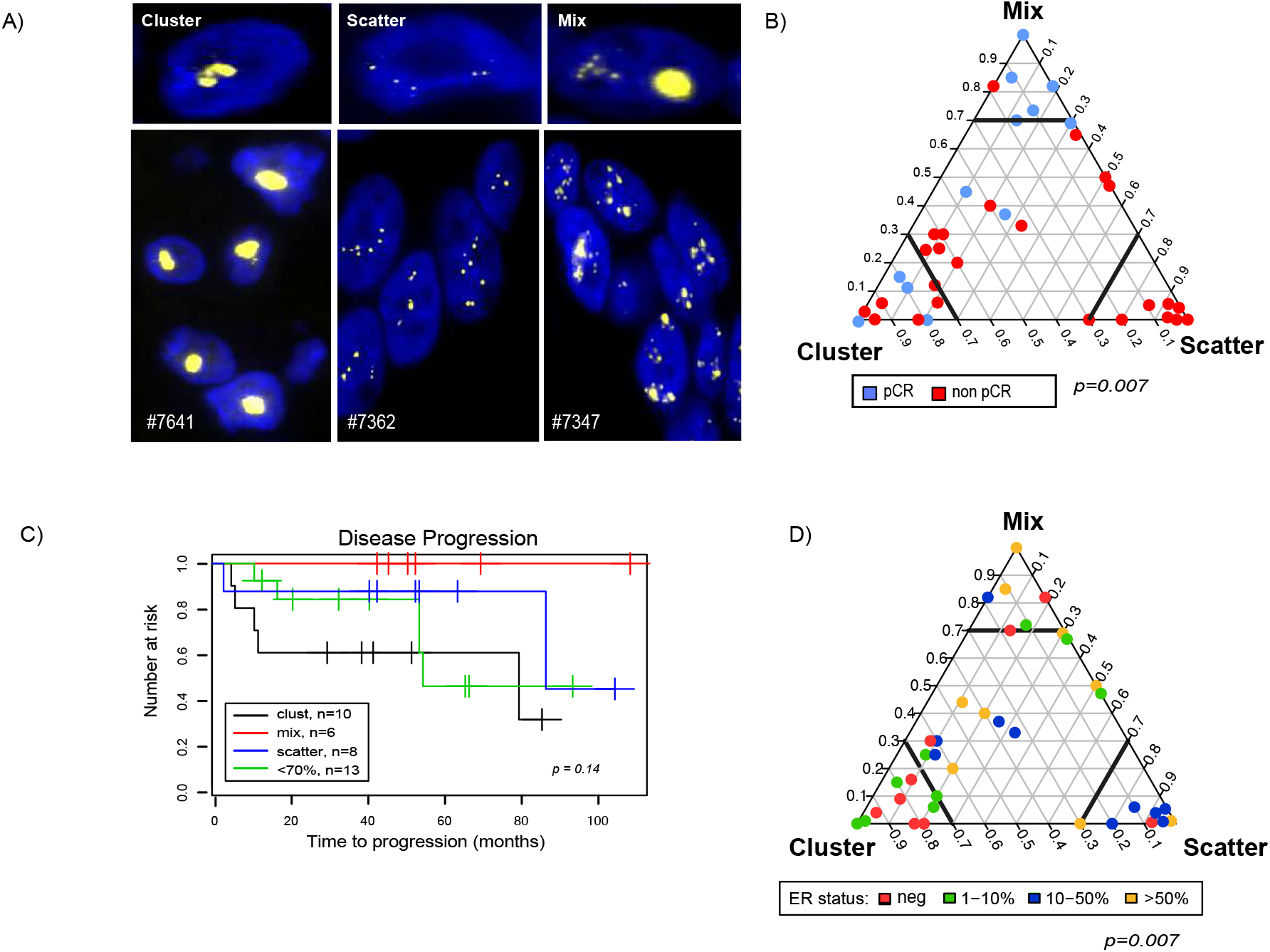
The spatial organization of the HER2 gene copies within the nuclei. A) Each cell was categorized as “cluster”, “scatter” and “mixed” based on the spatial organization of the *HER2* gene within the nuclei. B) The spatial organization for the HER2 CN for the pre-treatment samples (n=37), in the triangle plot each corner represent homogenous cell population (100% of cells have one of the spatial patterns). Samples from patients with complete response are colored in blue and from patients with non-complete response are colored in red. C) Kaplan-Meyer curve for time to disease progression for the categorized spatial organization “cluster”, “mix”, “scatter” and the “<70%” groups. D) The spatial organization for the pre-treatment samples where samples are colored by ER expression level (percentage of positive cells). ER negative samples are colored in red, ER low (1-10%) colored in green, ER intermediate (10-50%) colored in blue and ER high (>50% colored in yellow).

### 3.3 Predicting disease progression by measurements of clonal shift during therapeutic intervention

As patients with more heterogeneous tumors (reflected both by ER status and by cellular subclones displaying different types of *HER2* CN) had a higher risk of relapse, we next studied the population dynamics, i.e. which cell types responded or not to therapy and whether dynamics during therapy can reveal patients with better prognosis or not. We assessed change in tumor composition in 20 patients who did not achieve complete pathological response. To objectively address the dynamics of cell populations during neoadjuvant treatment, we calculated changes in fractions of the predefined cell types (phenotypic and *HER2* CN and the combined phenotypic/*HER2* CN cell types) before and after therapy using the Kullback-Leibler (K-L) divergence index. Figure 6A illustrates the change in *HER2* CN cell types (delta calculated by comparing fractions before and after therapy) sorted according to decreasing K-L index. Patients with low K-L index had a significant increased risk of breast cancer related death compared to patients with high K-L index, indicating that patients with smaller changes in subpopulations of cells during treatment actually have worse long-term outcome (Figure 6B, p=0.035). There was no correlation to any other clinico-pathological parameters, including degree of pathological response (Supplemental Table 6). Figure 6C shows IFISH images (*HER2* CN) from samples taken before and after therapy for two patients. Patient #7588 who did not have a progression of the disease showed a decrease in the fractions of cells with *HER2*amp, while patient #7435 who developed progression of the disease did not show any changes in the *HER2* CN cell types during therapy. In contrast, there was neither any association between patient outcomes with phenotypic changes nor with combined phenotypic/*HER2* changes based on the K-L index (Supplemental Figure 7A-B).

With regard to the individual markers analyzed, we did not observe any significant changes in the global levels of *HER2* and cent17 CN level, nor in the HER2 and ER protein intensity in tumors after neoadjuvant treatment (Supplemental Figure 7C). In particular we did not observe a significant difference between patients with a high shift of phenotype or combined phenotypic/*HER2* CN status compared to those with a low shift with regard to survival of disease or outcome.

**Figure 6:**
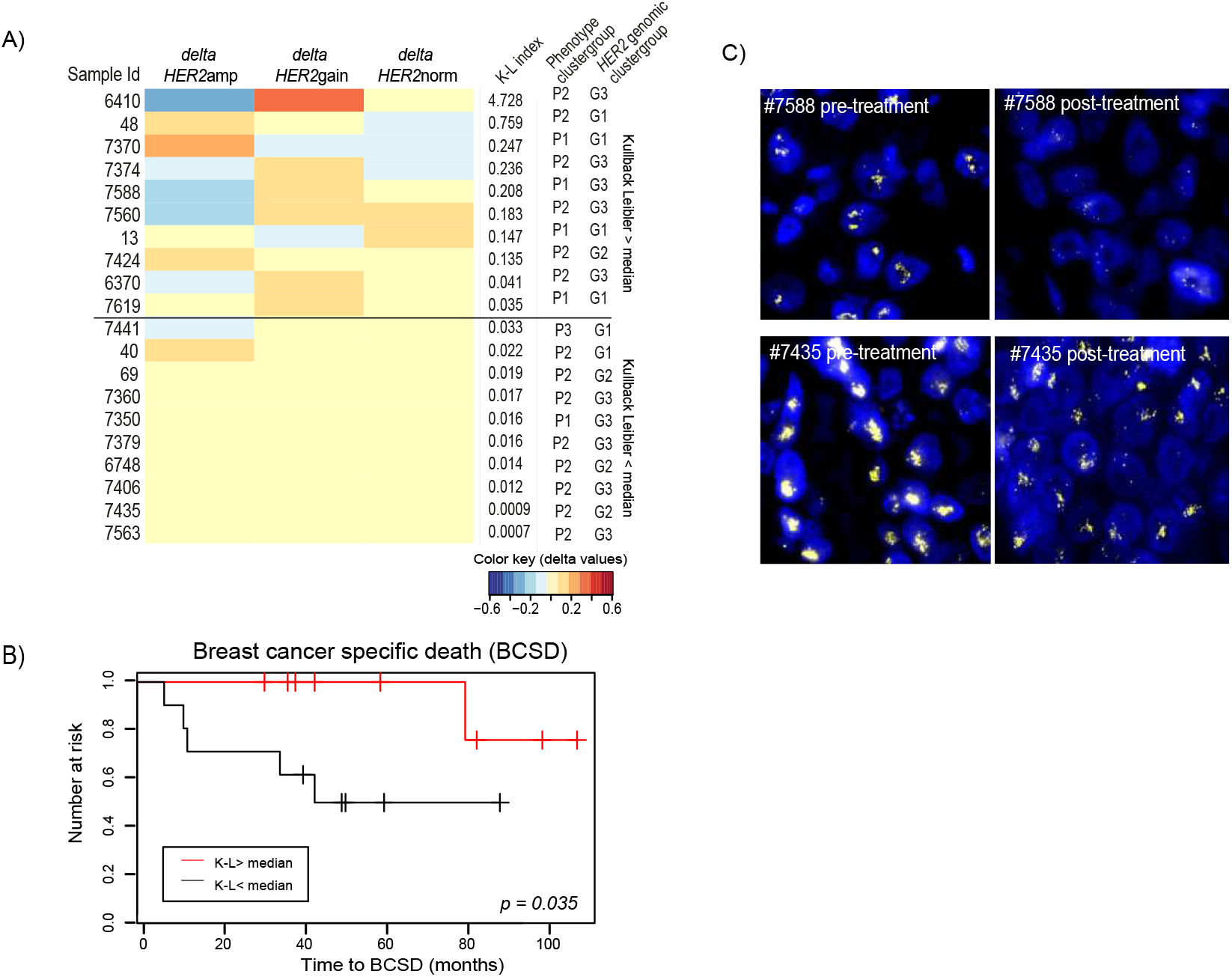
Tumor evolution during neo-adjuvant treatment. A) The Kullback-Leibler diversity index (K-L index) was calculated reflecting changes in cells with different levels of *HER2* CN during therapy. The samples were sorted from high to low K-L index, and the changes of the different cell typed from pre- to post-treatment are visualized by the delta values. To the right is the K-L index value and the genotypic and phenotypic cluster group for each patient. B) Example images from pre- and post-treatment biopsies from one patient with high K-L index (patient #7588) and from a patient with low K-L index (patient #7435). B) A significant increase in risk for death of breast cancer were seen for patients with low versus high K-L index (p = 0.035).

### 3.4 Diversity in primary tumor versus metastasis

Sampling of tumor metastases was not included in the study protocol, but tissue biopsies from distant metastases were available from three of the patients (two patients with non-complete response and one patient with complete response to therapy). IFISH images of biopsies from three time points (pre- and post- treatment and later distant metastasis) of two of the patients are shown in Figure 7A-F. Patient #7435 (Figure 7A-C) had a primary tumor dominated by HER2+/ER-cells with *HER2* CN amplification. After neoadjuvant treatment we found an increase in cells with HER2+/ER+ phenotype. Interestingly, the biopsy from a metastasis showed the same cell phenotypes as the pre-treatment tumor. There was no evidence of clonal shift as the samples from all three time-points were dominated by cells with *HER2* CN amplification (Figure 7G). In contrast, the tumor from patient #7360 (Figure 7D-F) had prior to treatment mainly HER2+/ER-cells, but the biopsy after treatment and from the metastasis revealed a small fraction of HER2-/ER-cells. There was only a minimal change in the fraction of cells with *HER2* CN amplification (Figure 7H). We also investigated the spatial organization of the *HER2* CN at the three time points, and both samples had a more similar spatial pattern for the *HER2* CN for the pre-treatment and metastatic lesion in contrast to the post-treatment biopsy, but the changes were only minor (Figure 7I and 7J).

**Figure 7:**
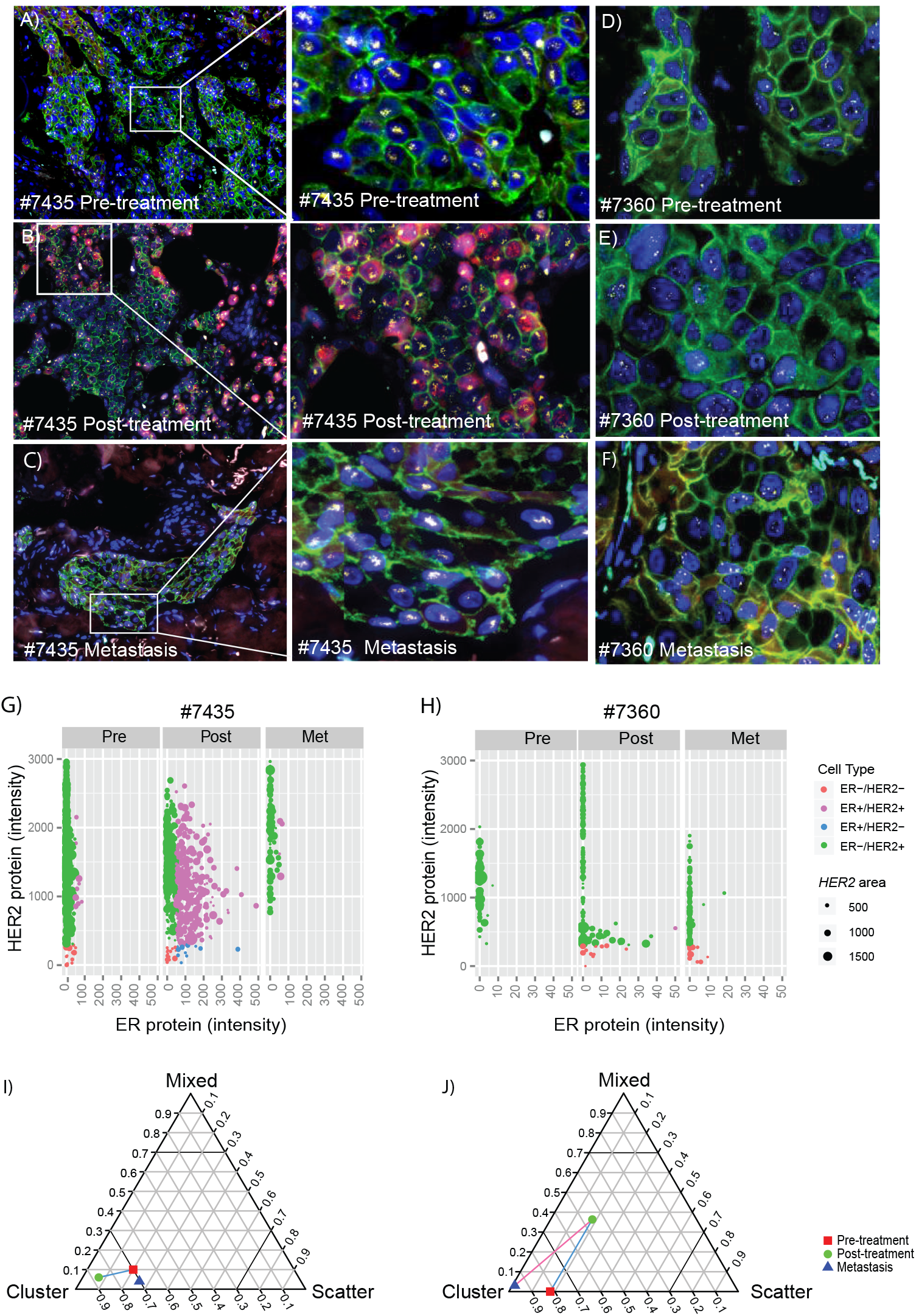
Intra-tumor heterogeneity during disease progression. IFISH images from biopsies from patient #7435 (with a magnified area to the right): A) pre-treatment biopsy, B) post-treatment biopsy and C) biopsy from a metastasis. Equally from patient #7360: D) pre-treatment biopsy, E) post-treatment biopsy and F) biopsy from metastasis (Dapi=blue, HER2=green, ER=red, *HER2*=yellow and cent17=cyan). The phenotype and *HER2* CN level for all tumor cells analyzed from each of the three biopsies are plotted in the diagram G) patient #7435 and patient #7360 (colored due to their phenotypic cell type and the size of the spot reflect the *HER2* copy number level). Spatial organization of the *HER2* gene visualized in a triangle for the pre- (red square), post- (green circle) and metastatic- (blue triangle) sample from patient #7435 (I) and patient #7360 (J).

## 4 Discussion

Analysis of tumor samples taken from patients during neoadjuvant treatment is extremely useful for studying the clinical impact of tumor cell diversity. The significance of intra-tumor heterogeneity for treatment response can be measured by comparing molecular features of tumor cells from pre- and post-treatment biopsies. As *in situ* methods only allow us to measure a small number of markers, we chose the clinically most important biomarkers, namely ER (protein) and HER2 (protein and gene copy number). Even with so few biomarkers, the combined IFISH technique revealed a high diversity both between tumors but also within tumors (i.e. cell-to-cell variation). It is known that tumors classified as HER2+ by immunohistochemistry (i.e. 3+) can have different levels of *HER2* amplification by ISH techniques. Our work supports this observation but also provide a higher resolution as all markers are studied simultaneously in thousands of individual cells. We found remarkable diversity, both with regard to the expression of ER and HER2 protein as well as for *HER2* CN on single cell level (Supplemental Figure 1, 3 and 4). It was intriguing to find some tumors with a linear correlation between the two proteins and/or between protein and *HER2* CN, while others were not linear. This prompted us to classify each cell into phenotypic and genomic predefined categories. By performing three separate clustering analyses we found several interesting features characterizing the tumors of patients with a higher risk for disease progression and/or breast cancer related death: (i) high expression of HER2 but low or intermediate number of ER+ cells (P2 in Figure 3), (ii) a mixture of cells with different *HER2* CN levels (G3 in Figure 4) and (iii) a mixture of cells with different *HER2* CN levels with low number of ER+ cells (PG3 in Supplemental Figure 5). Combined, these findings indicate that patients with tumors dominated by *HER2* amplified cells and with homogenous ER expression (either negative or positive) have a good long-term prognosis. It also indicates the importance of addressing not only the heterogeneity of *HER2* CN but also the variation in ER expression in HER2+ breast carcinomas. In the work by Ferrari et al. (Ferrari et al. 2016), HER2+ tumors were split into four groups based on gene expression patterns, and the level of ER expression varied between them. Although the study did not address intra-tumor heterogeneity, it clearly showed that a subgroup of HER2+ carcinomas was composed of ER negative tumors, one subgroup of highly ER positive and two subgroups of tumors with more intermediate ER levels. It will be of interest to see the follow-up studies of this cohort with outcome data as well. In a recent study, approximately 30% of patients with neoadjuvant treated HER2+ tumors (chemotherapy and HER2 targeted treatment) achieved pathological complete response (pCR), but this fraction was lower for patients with HER2+ and ER+ tumors, but the level of ER positivity was not addressed (Cortazar et al. 2014). In a study by Rodmond et al., patients with ER+ tumors had a lower response rate to treatment, but this seems to be mainly restricted to those with tumors having less than 50% ER positive tumor cells (Romond et al. 2005). These findings are in line with ours; patients with heterogeneous ER expression had a tendency towards a reduced long-term survival (Figure 3). Carey et al. recently published results from the CALGB40601 trial, which also shows that local response varies between ER+ and ER-subtypes of HER2+ breast cancer (Carey JCO 2016). We found no evidence that the HER2 protein intensity level has impact on local response, which is in line with the observation by Zabalgo et al. (Zabalgo Ann of Onc 2013) but contradicts the CALGB 40601 trial which found gene expression levels of both ER and HER2 to be correlated with pCR rates (Carey JCO 2016).

In our study, we find *HER2* CN level to be of clinical importance as the level in pre-treatment samples was significantly higher in tumors from responders compared to non-responders. This is in line with previous studies showing high levels of *HER2* amplification to be associated with pathological complete response (pCR) (Arnould et al. 2007)(Guiu et al. 2010) although *HER2* CN level could not predict long-term disease progression or survival. This is supported by studies of anti-HER2 treatment in adjuvant setting where *HER2* CN level has shown no or negative correlation with disease free survival (Xu et al. 2016). As mentioned previously, *HER2* CN heterogeneity seems to have impact on prognosis in our study. We found tumors with heterogeneous composition with regard to *HER2* CN level to have higher risk of relapse and breast cancer specific death (patients in G3 group in Figure 4). Some studies indicate the same result in less advanced stage of the disease; in a study of adjuvant treated HER2+ breast cancer, Seol et al. found regional heterogeneity in *HER2* CN to predict a worse survival (Seol et al. 2012). The study by Lee et al. also found patients with both regional and genomic heterogeneity of *HER2* amplification to have decreased disease free survival, but neither of these two study cohorts had uniform treatment regimens (Lee et al. 2014; Seol et al. 2012). Korozumi et al. studied variation in both *HER2* copy number and HER2 protein expression within tumors using a semi-objective analysis (with visual scoring) and found that regional variation of *HER2* CN reflected a worse prognosis particularly in ER negative disease (Korozumi 2016). Unfortunately, these patients had not received anti-HER2 therapy, so neither the predictive value nor the impact of dynamics during therapy could be addressed.

One of the most striking findings in our study was the large number of tumors exhibiting intra-tumor variation with regard to *HER2* CN levels. As copy number alterations are inherited in daughter cells, we believe these populations to reflect true sub-clones that have undergone different paths of evolution. The cluster analysis based on *HER2* CN levels showed that patients with tumors dominated by cells with amplified *HER2* gene had a significant better survival compared to the patients with a more heterogeneous *HER2* amplification levels (Figure 4). Patients in the latter group (Cluster G3 in Figure 4) had tumors with a mixed cellular composition. These patients had a significant shorter time to progression of the disease and fewer long-term survivors. We suggest that patients belonging to cluster group G3 represents cases similar to those described by Ballard et al. as “non-classical” *HER2* FISH results (Ballard et al. 2017).

Changes in ER and HER2 status is observed for some cases during neoadjuvant treatment, and this change seems to affect protein expression (i.e. phenotype) more than *HER2* copy numbers (Van de Ven et al. 2011). However, studies of genomic and phenotypic intra-tumor heterogeneity of HER2+ breast carcinomas and their impact on treatment resistance have been scarce. A recent work studying HER2+ tumors at single cell level found overexpression of *BRF2* and *DSN1* genomic driver events in HER2 negative cells (Ng et al. 2015). This indicates a presence of subpopulations that can explain treatment resistance. It has also been shown that that important genetic driver events such as *PIK3CA* mutation and *HER2* gene amplification is not always present within the same cell (Janiszewska et al. 2015). As minor subclones might need time to proliferate and progress (by clonal selection), this could explain why we find heterogeneous tumors to have a significant increased risk for disease progression regardless of the initial local response. When comparing the intra tumor heterogeneity before and after treatment, we were surprised to find that patients in the group with no changes in the cellular composition had an increased risk for later progression of the disease. One explanation for this finding could be that none of the tumor sub-clones were affected by the treatment and probably reflecting tumors where *HER2* gene amplification is not the important driver. Another explanation could be treatment resistance due to ligand independent activation of HER2 (Yarden 2001) rather than selection of clones proliferating independently of HER2 activity. Interestingly, these tumors do not reflect the situation identified by Ng et al. where a HER2 negative subpopulation could be suspected to explain therapy resistance (Ng et al. 2015). Our study was unfortunately not suitable for Next Generation Sequencing NGS based identification of driver events in resistant subclones and more detailed explorative studies to identify alternative candidate drivers will be needed. Identification of distinct genomic alterations related to the cellular dynamics during treatment might provide clinicians with more therapy options for such patients.

Finally, the cases with samples from three time points showed intriguing results; the pre-treatment and metastatic lesion had a more similar spatial pattern for the *HER2* CN in contrast to the post-treatment biopsy (Figure 7I and 7J). One of the cases showed a major switch in phenotype (Figure 7A-C) but had a very low Kullback-Leibler index, reflecting minor influence of treatment on *HER2* CN cell types. The other case had only a minor phenotype change and the *HER2* CN cell types did not shift enough to be reflected by the Kullback-Leibler index. Although this is just case observation, it reflects breast cancer to be a disease that can evolve along different paths both with regard to phenotype and genomic/clonal composition.

An important challenge for estimating intra-tumor heterogeneity is the need for objective measurements of molecular biomarkers. Buckley et al. proposed a simple heterogeneity index for *HER2* CN heterogeneity, but this was based on visual counting of 20 cells (as defined by the CAP guidelines) by an observer (Buckley et al. 2016). To address potential observer bias and maximize the number of analyzed cells, we estimated heterogeneity by objective assessment of *HER2* CN of more than 13.000 cells using GoIFISH, an image analysis software that can omit artificial staining and specifically characterize tumor cells for further analysis. Still, tissue artifacts such as incomplete tumor cell nuclei due to sectioning can influence the results. We also used cluster analyses of the *fractions* of cell types within a tumor, thus the presence of some misclassified cells will not influence the results substantially. Finally, the visual categorization of intra-nuclear spatial distributions of the *HER2 amplicon* also reflected the presence of different types of genomic disruptions and amplification mechanisms, representing a different way of assessing clonal heterogeneity. Here we analyzed fewer cells per sample (100 cells), but the finding is in line with other studies (by DNA sequencing) showing that *HER2* gene amplifications can be a result of different types of DNA rearrangement mechanisms (Morganella et al. 2016). This cohort does not have tumor material suitable for NGS analyses of this kind, but this is important to address in suitable sample collections.

This study is based on a neoadjuvant observational trial, comprising of HER2+ patients for which matched primary, post-treatment and in some cases metastatic samples were available for analysis. The strength of this cohort lies in the strict inclusion criteria and consistency in terms of treatment regimens, allowing us to make direct comparisons between patient samples and track the cellular dynamics throughout the treatment process. Although this study could benefit from an increased patient sample size and sufficient patient material to conduct DNA sequencing analysis, this observational cohort has nonetheless offered an insight on the wide biological specter within HER2+breast carcinomas and in particular the negative association between *HER2* CN intra-tumoral heterogeneity and patient outcome.

## 5 Conclusion

This is to our knowledge the first study of breast cancer revealing cellular heterogeneity with regard to HER2 expression, *HER2* copy number and ER expression in analyzing a substantial number of cells from neoadjuvant treated HER2+ breast cancer patients. HER2+ disease is highly heterogeneous both between and within tumors. The heterogeneity of ER expression as well as *HER2* copy number variation seems to have impact on disease progression and survival. Additionally, tumors with preserved level of heterogeneity during therapy with regard to *HER2* CN types (i.e. cell-type composition before and after therapy) had a poor prognosis. The study shows the importance of assessing cell-to-cell variation both prior to treatment but also during treatment, and measuring shifts in cell populations has a potential when it comes to predicting therapy response. It also shows the importance of having an objective analysis of multiple markers in a high number of cells facilitated by automatized image analysis. The challenge now is not only to validate the clinical impact of molecular subtypes within HER2+ breast cancer patients, but also to address the cellular variation within the tumors in more depth.

## Acknowledgements

We thank the hospitals in Buskerud, Telemark, Vestfold, Østfold, Bodø, Innlandet, AHUS, Ullevål and the Norwegian Radiumhospital for retrieving and sending us the tissue blocks from the patients included in this study.

IHR and HGR are grateful for funding from Health Region South-East, Norway and from The Norwegian Cancer Society. IHR, HGR and ALBD are also grateful for support for accommodation in Cambridge, UK from Radiumhospitalets legater. FM would like to acknowledge the support of The University of Cambridge, Cancer Research UK and C K Hutchison Holdings Limited. Parts of this work were funded by CRUK core grant C14303/A17197 and A19274.

## Author contributions

HGR, ÅH, ALB-D and IHR conceived the study. Trial design and clinical data by AS and ÅH. IHR, VA, KP and HGR conducted and/or designed all wet-lab experiments and tissue analyses. IHR, AT, FM, DN, and OCL performed and/or developed image analyses, bioinformatical analyses and statistical analyses. HGR, FM, IHR, AT, ÅH, AS, ALB-D, KP, VA, DN and OCL participated in the combined analyses and wrote the manuscript.

## Supporting information

All computational scripts used are available in: Supplementary_Sweave.Rmd

